# Early, color-specific neural responses to object color knowledge

**DOI:** 10.1101/2021.02.01.429104

**Authors:** Talia L. Retter, Yi Gao, Fang Jiang, Bruno Rossion, Michael A. Webster

**Affiliations:** Department of Psychology, Center for Integrative Neuroscience, University of Nevada, Reno (USA); Psychological Sciences Research Institute, Institute of Neuroscience, UCLouvain (Belgium); Department of Behavioural and Cognitive Sciences, Institute of Cognitive Science & Assessment, University of Luxembourg (Luxembourg); Université de Lorraine, CNRS, CRAN, F-54000, Nancy (France); Université de Lorraine, CHRU-Nancy, Service de Neurologie, F-54000 Nancy (France)

**Keywords:** memory, visual cortex, shape, frequency tagging, EEG

## Abstract

Some familiar objects are associated with specific colors, e.g., rubber ducks with yellow. Whether and at what stage neural responses occur to these color associations remain open questions. We tested for frequency-tagged electroencephalogram (EEG) responses to periodic presentations of yellow-associated objects, shown among sequences of non-periodic blue-, red-, and green-associated objects. Both color and grayscale versions of the objects elicited yellow-specific responses, indicating an automatic activation of color knowledge from object shape. Follow-up experiments replicated these effects with green-specific responses, and demonstrated modulated responses for incongruent color-object associations. Importantly, the onset of color-specific responses was as early to grayscale as actually colored stimuli (before 100 ms), the latter additionally eliciting a conventional later response (approximately 140-230 ms) to actual stimulus color. This suggests that the neural representation of familiar objects includes both diagnostic shape and color properties, such that shape can elicit associated color-specific responses before actual color-specific responses occur.

## Introduction

Objects typically are specifically defined by their shape, and thus shape information is often sufficient for basic-level visual object recognition (Grossberg and Mingolla, 1985; Biedermann 1987; Biederman & Ju, 1988). While surface attributes, such as texture and color, are often not object-specific, they nevertheless contribute substantially to object recognition (Marr & Nishihara, 1978; Tanaka, Weiskopf, & Williams, 2001). In part, color may assist in object recognition by providing cues to image segmentation and thus shape: for example, color can be more reliable than luminance for defining object boundaries, since luminance variations often also arise from shading (Kingdom, Beauce & Hunter, 2004), and color is among the most powerful cues for perceptually grouping and spatially organizing the visual stimulus (Wolfe & Horowitz, 2004). Yet color also provides object-diagnostic information, for even clearly segmented objects shown in color are typically recognized faster and more accurately than when shown in grayscale (Price & Humphreys, 1987; Wurm et al., 1993; Humphrey et al., 1994; Tanaka & Presnell, 1999; Nagai & Yokosawa, 2003; Therriault, Yaxley, & Zwaan, 2009; Rossion & Pourtois, 2004; Bramao et al., 2011; Hagen et al., 2014).

The contribution of color to object recognition is not equal for all objects. For example, color contributes more towards the recognition of structurally similar objects (e.g., some fruits and vegetables) or when an object exemplar has an atypical or degraded shape (Markoff, 1972; Price & Humphreys, 1989; Wurm et al., 1993; Joseph & Proffitt, 1996; Tanaka & Presnell, 1999; Liebe et al., 2009). Yet, the objects benefiting the most from the presence of color are those for which familiar objects have a learned association with a specific color, so-called “color-diagnostic” objects (e.g., Helmholtz, 1867; Joseph & Proffitt, 1996; Tanaka & Presnell, 1999; Naor-Raz, Tarr, & Kersten, 2003; Rossion & Pourtois, 2004; for a meta-analysis: Bramao et al., 2011). Such color-associated objects are present across a variety of object types, including natural, artificial, living, non-living, and animated: e.g., blueberries (blue), lobsters (red), US dollar bills (green), and Pikachus (yellow; for further examples of color-associated objects, see Appendix B of Joseph & Proffitt, 1996; Table 1 of Tanaka & Presnell, 1999; Appendix 1 of Naor-Raz, Tarr, & Kersten, 2003; Appendix A of Nagai & Yokosawa, 2003; Appendix 5 of Rossion & Pourtois, 2004; Fig. 3 of Witzel et al., 2011). When such color-associated objects are presented in incongruent colors, i.e., any color other than their typical color, object recognition performance is lower in terms of accuracy and/or response time relative to objects presented in congruent color and, to a lesser extent, in grayscale (Price & Humphreys, 1987; Joseph & Proffitt, 1996; Tanaka & Presnell, 1999; Therriault, Yaxley, & Zwaan, 2009; Hagen et al., 2014). For these objects, the specific color/object association allows color to provide a diagnostic cue for object recognition. Moreover, it is possible that this association allows object shape to provide a cue for object color: color naming is less accurate and/or delayed for incongruent relative to congruent color-associated objects (Bruner & Postman, 1949; Ratner & McCarthy, 1990; Naor-Raz, Tarr, & Kerseten, 2003).

These observations suggest that the recognition of color-specific objects by shape alone automatically evokes their color associations. Indeed, the perception of color-specific objects may be drawn towards the associated color of the object, i.e., the “memory color”, particularly when the presented color is categorically ambiguous, e.g., yellow-orange, or presented in the context of challenging viewing conditions (e.g., Duncker, 1939; Bruner & Postman, 1949; Mitterer & de Ruiter, 2008; Vandenbroucke et al., 2016). Such color association effects have been further demonstrated in work suggesting that grayscale object images can appear subtly tinged with their characteristic hue (Hansen et al., 2006; Olkkonen, Hansen, & Gegenfurtner, 2008; Witzel et al., 2011), and that chromatic afterimages of objects appear more vivid when corresponding to their typically associated colors (Lupyan, 2015).

Functional neuroimaging studies have reported that color-associated grayscale objects activate visual regions of the human brain associated with color perception, including in the fusiform gyrus (Martin et al., 1995; Simmons et al., 2007; Slotnik, 2009). More recently, these findings have inspired the successful decoding of the associated color of grayscale color-specific images with functional neuroimaging in visual regions or whole-brain analyses, even when subjects performed orthogonal, i.e., non-color-related, tasks (Bannert & Bartels, 2013; Vandenbroucke et al., 2016; Teichmann et al., 2018; 2019). However, some of these studies have used only a few objects per color category (e.g., two in Bannert & Bartels, 2013), and have reported divergent results in terms of the cortical areas responding to memory color (V1: Bannert & Bartels, 2013; V3, V4, VOI, LOC, and prefrontal areas: Vandenbroucke et al., 2016). Nevertheless, in light of these findings, it appears possible that color-specific brain responses are automatically activated by the visual shape of the color-associated objects. However, how shape information might elicit an associated color response, and how that response differs from the response to an actual color stimulus, remain unknown.

These issues directly refer to the critical question of the stage(s) of visual object processing/representation at which color and shape components are related. On one hand, a late role of color (e.g., Marr & Nishihara, 1978; Biedermann 1987; Proverbio et al., 2004; Therriault, Yaxley, & Zwaan, 2009; Proberbio et al., 2004; Teichmann et al., 2019), is proposed in a hierarchical view of visual perception, In this view, color processing is only influenced by shape-based expectations at later stages of visual processing, i.e., following feedback from high-level visual areas. This implies that chromatic and spatial representations are relatively independent at early visual stages, which has been supported by some lines of evidence. For example, color or shape differences (e.g., in orientation or size) are salient cues supporting rapid visual search, yet an arbitrary color-shape conjunction requires attention to be detected (e.g., Treisman, 1982; Wolfe, Cave & Franzel, 1989). Similarly, some aspects of spatial vision (e.g., acuity and disparity processing) and motion perception can become degraded for images defined only by chromatic contrast (Cavanagh, Tyler & Favreau, 1984; Mullen, 1985; Shevell and Kingdom, 2008), and color tends to be more labile and fill-in between boundaries defined by spatial variations in luminance (Pinna, Brelstaff & Spillmann, 2001; van Lier, Vergeer & Anstis, 2009), results which have fueled substantial interest in the modularity of visual processing (e.g. Zeki & Shipp, 1988; Livingstone & Hubel, 1984).

On the other hand, an early role for associated color (e.g., Price & Humphreys, 1989; Naor-Raz, Tarr, & Kersten, 2003; Rossion & Pourtois, 2004; Lu et al., 2010; Hagen et al., 2014; Teichmann et al., 2018) is present in a framework in which object color knowledge, through past experience, constrains, and perhaps even generates, color perception (e.g., Helmholtz, 1867; Gregory, 1966; Thompson, 1995; Lotto & Purves, 2002). This view necessitates that object color and shape are rapidly, automatically integrated in object recognition. There is evidence for such an interaction in early visual processing: for example, cells in the primary visual cortex are often tuned to both spatial and chromatic information (Thorell, De Valois & Albrecht, 1984; Johnson, Hawken & Shapley, 2001) and can be selective to conjunctions of color and form (Seymour et al., 2010). Psychophysically, adaptation and masking are selective for both the shape and color of the stimuli (McCollough, 1965; Bradley, Switkes & De Valois, 1988; Clifford et al., 2003), and chromatic contrast can support many spatial discriminations (Webster, De Valois & Switkes, 1990; Krauskopf and Farell, 1991). Evidence for both early and late roles of color, and how it relates to shape in visual object processing, do not limit an influence of color to any stage of visual processing (Johnson & Mullen, 2016).

The few electro/magnetoencephalography (EEG and MEG, respectively) findings on the timing of color in object recognition are inconsistent. Teichmann et al. (2018) reported earlier color-specific responses to colored than grayscale objects, based on the success of a multivariate pattern-analysis classifier trained on colored objects with full-brain MEG, but did not observe this pattern in a later study (2019). Other studies have not reported significant latency differences between event-related potentials (ERPs) evoked by color, grayscale, or incongruently colored objects (Proverbio et al., 2004; Lu et al., 2010; Bramao et al., 2012b; Lloyd-Jones et al., 2012). In these studies, the latency of color-associated effects has been inferred from the onset of amplitude differences across color *vs*. grayscale or congruent *vs*. incongruent color objects. Such differences were reported at the earliest components in some studies (Lu et al., 2010: N1 and later components decreased for congruent color *vs*. grayscale/incongruent color; Bramao et al., 2012b: P1 and N1 increased for color *vs*. grayscale), but only at later components in other studies (Provebio et al., 2004: N2 increased when attending to color for congruent *vs*. incongruent shapes; Lloyd-Jones et al., 2012: P2 and P3 decreased for congruent *vs*. incongruent color; Bramao et al., 2012b: N400 increased for diagnostic vs. non-diagnostic color). A common limitation of these studies is that responses to different object color categories were averaged together, despite evidence that the spatiotemporal dynamics of EEG responses to different colors can differ substantially (e.g., Regan, 1966; Riggs & Sternheim, 1968; Allison et al., 1993; Anllo-Vento, Luck & Hillyard, 1998; Retter et al., 2020).

In the present study, we address the unresolved issue of the processing/representational stage(s) at which color and shape components of a visual object are jointly encoded by using a sensitive and objective frequency-tagging EEG approach (Norcia et al., 2015). First, we quantify the color-specific neural responses to familiar, color-associated objects, using frequency-tagging to isolate these responses from the generic responses to visual stimulation, and from the effect of color on these generic responses. Second, using grayscale versions of these objects, we quantify the contribution of any associated color signal to the color-specific neural response. Finally, we compare the temporal dynamics of the neural responses to actual *vs*. implied color-specific stimuli. Our findings indicate that while there is an effect of color on visual object stimulation responses, color-associated object shape alone can evoke substantial color-specific neural responses. Moreover, the implied color response emerges as early as the corresponding responses for actual colored objects, with the neural responses for the two types of stimuli diverging only later in time.

## Methods

### Participants

Sample size per experiment was determined relative to previous within-samples, frequency-tagging EEG studies concerning color perception, for which the sensitivity and reliability, even at the individual level, is high (e.g., 14 participants, medium effect sizes for color differences, over 80% of individuals with significant responses: Retter et al., 2020; Or, Retter & Rossion, 2019; see also the review of Norcia et al., 2015). Sixteen participants took part in Experiment 1 (targeting yellow-associated object responses), who aged from 18 - 35 years old (M = 24.5 years; SD = 5.27 years); eleven identified as female, and five as male; thirteen as right-, and three as left-handed. Sixteen participants also took part in Experiments 2 and 3, in a single testing session. These latter participants were aged 20 - 30 years old (M = 24.6 years; SD = 3.67 years); eleven identified as female, and five as male; fourteen as right- and one as left-handed, and one as ambidextrous. All reported normal or corrected-to-normal visual acuity, as well as normal color vision. All participants gave signed, informed consent prior to participation. Each experiment was performed only once, and no participants were excluded from the analyses. The study was conducted in accordance with the Code of Ethics of the World Medical Association (Declaration of Helsinki), and approved by the University of Nevada, Reno’s Institutional Review Board.

### Stimuli

Objects with a strongly associated color were chosen with considerations for color category and object type. Ultimately, 24 object images were selected, balanced for color category (red, green, blue, and yellow) and object type (fruit/vegetable, cartoon character, manmade, and animal), and controlled for low-level attributes (**Fig. 1a**).

**Figure 1.**
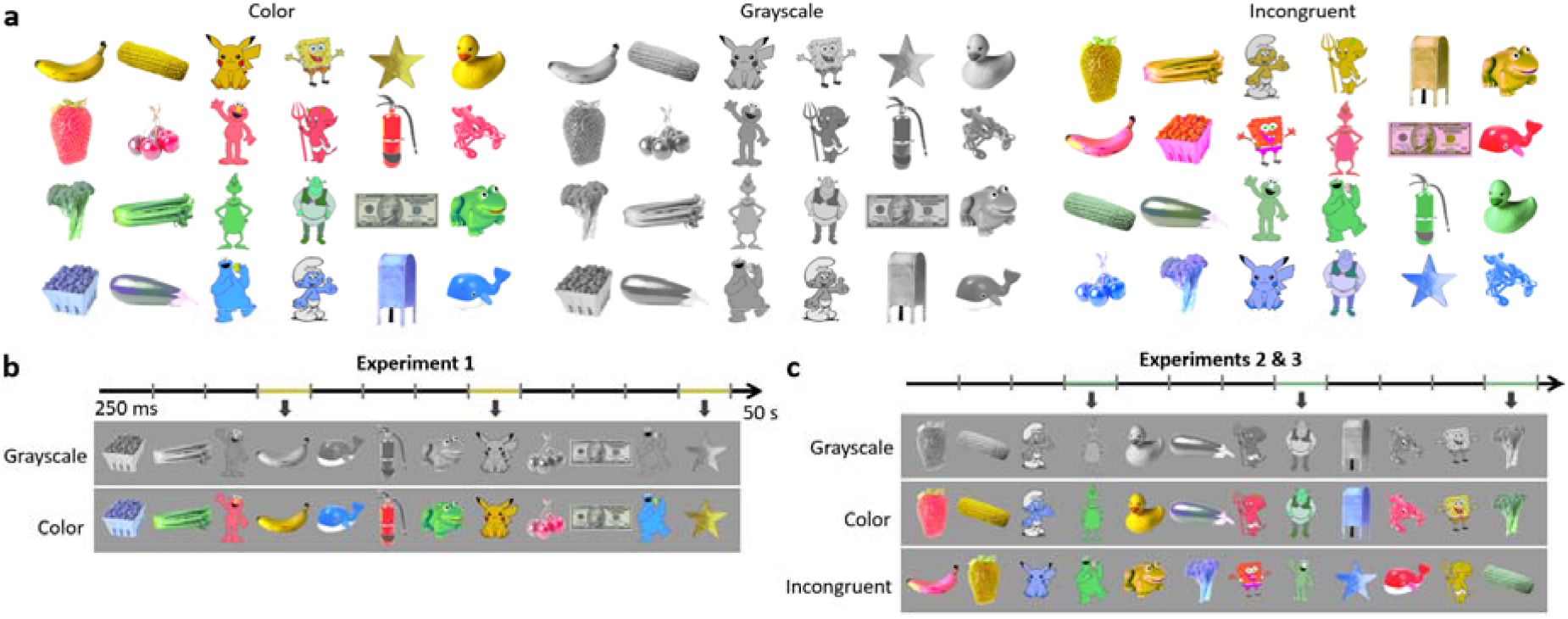
**a)** The 24 color-associated stimuli used in the experiments, shown as appearing in diagnostic color (left), grayscale (middle), and incongruent color (right) conditions. **b)** The trial design of Experiment 1, targeting yellow-associated objects, is depicted for each the grayscale and color conditions. Stimuli were presented every 250 ms (at 4 Hz) in 50 s sequences, throughout which a yellow-associated object appeared every 1 s (1 Hz), i.e., as every fourth image. The order of the green-, red-, and blue-associated images was fully randomized within every sequence for each participant. **c**) The trial design of Experiments 2 and 3, targeting green-associated object responses. Experiment 3 consisted of an incongruent condition, in which targeted non-green-associated objects appeared in green.

With regard to color association strength (“color diagnosticity”), we first selected images of 40 candidate objects, with reference to previous studies (Tanaka & Presnell, 1999; Naor-Raz, Tarr, & Kersten, 2003; Witzel et al., 2011). In an informal survey, we presented these images in grayscale to a room of 110 undergraduate students. Participants were asked “what color do you think the item is?”, and instructed to fill in a sheet to indicate their response from the following list: red, orange, yellow, green, blue, purple, gray, and don’t know. Color association was assessed by the percent agreement across participants for the most frequent color selected. Objects with the highest color association ratings were selected while maintaining balance in color category and object type, resulting in a minimum of 93% color naming agreement on average within each color category (range: 93 to 96%).

The resulting stimuli of each color category consisted of two fruits/vegetables, two cartoon characters, one manmade object, and one animal. These include for red: strawberry, cherries, Elmo, Devil, fire extinguisher, and lobster; yellow: corn, banana, Pikachu, SpongeBob, star sticker, and rubber duck; green: broccoli, celery, Grinch, Shrek, dollar bill, and frog; blue: blueberries, eggplant, Cookie Monster, Smurf, mailbox, and whale. Note that the eggplant was actually named as purple, but was nevertheless included in the blue set to match for object type. The object images were coarsely selected for similarity in visual appearance: for example, the cartoon characters were restricted to two-dimensional renderings with similar postures, and the animal was consistently a toy version.

To control for low-level attributes, the images of these objects were first isolated from their background, cropped to their external edges, and resized to a common rectangular area. A grayscale set of these images was created with custom software, and both the colored and grayscale set were equalized in terms of mean luminance and root mean-squared luminance contrast. The colors for the different objects were adjusted to coincide with different hue angles relative to neutral gray (CIE 1931 x,y = 0.310, 0.316) within a version of the MacLeod-Boynton chromaticity diagram scaled to roughly equate threshold sensitivity along the cardinal axes (Winkler et al., 2015). For the different color categories, the mean hue angle in the space was set to: 355° (red), 315° (yellow), 205° (green), and 135° (blue), with all images’ mean hues restricted to a 10° range within each category, in order of the mean hue rankings of the original images. A set of incongruent color stimuli was made for Experiment 2, by assigning a non-associated color category to each image, while maintaining balance across object type.

To test whether additional image attributes might vary across object color categories, the grayscale objects were also examined in terms of additional (higher-order) image statistics, namely in terms of spatial frequency content (using scripts from Torralba & Oliva, 2003; Bainbridge & Oliva, 2015), global contrast factor (Matkovic et al., 2005) and gist (spatial envelope; Oliva & Torralba, 2001). No significant differences were found (see **Supplemental Material**), although it remains a possibility that the stimuli have some mean differences by color category (e.g., in curvature, local contrast, etc.). Note that the neural responses to grayscale objects may be expected to be particularly uniform across object examplars within participants, since color memories have been shown to be biased to prototypical color category membership (for objects: Van Gulick & Tarr, 2010; for color patches: Boynton et al., 1989; Uchikawa & Shinoda, 1996; Bae et al., 2015; see also Bartleson, 1960).

### Experimental Design

There were two conditions in Experiment 1, targeting yellow-specific responses: *grayscale* and *color* images. In the former, all images were presented in grayscale, and in the latter, the same images were presented in color. In Experiment 2, these two conditions were replicated while targeting green-specific responses. In Experiment 3, a novel condition targeting green-specific responses was tested: *incongruent* images, in which the color of each of the object images was replaced with a non-associated color (**Fig. 1a right**). Participants were presented with four 50-s sequence repetitions for each condition, leading to a total of 3.3 minutes of recording per condition (6.7 minutes of recording in Experiment 1; 10 minutes in Experiment 2). The conditions were presented in blocked order, counter-balanced across participants. In Experiments 2 and 3, the grayscale condition was always presented first, to prevent an influence on the object’s associated color from having first seen the incongruently colored images.

Throughout each sequence, stimuli were presented at a rate of 4 Hz, i.e., every 250 ms. With a 50% squarewave duty cycle, each image was displayed at full contrast for 125 ms, followed by 125 ms of the gray background between successive images. Crucially, in Experiment 1, yellow-associated images appeared every 1 s, at a rate of 1 Hz, within the sequence; in Experiment 2, green-associated images appeared at 1 Hz; in Experiment 3, incongruently green-colored objects appeared at 1 Hz. The order of presentation of the other, non-target color-associated objects was fully randomized within every sequence for every participant (**Fig. 1b**). Thus, responses to a consistent object-color association (i.e., yellow-selective responses in Experiment 1; green-selective responses in Experiment 2) were expected at 1 Hz and its harmonics (2, 3, 5 Hz, etc.), while responses to object presentation in general were expected at 4 Hz and its harmonics.

Participants could initiate trials at their own pace. Upon commencement of each trial, the testing sequence was preceded by 1 – 2 s of a fixation cross, in order to orient attention and decrease exact prediction of image onset, followed by 1 s of gradual stimulus contrast increase (fade-in), to avoid abrupt eye movements. Each sequence was followed by 1 s of fade-out and 1 – 2 s of the fixation cross, with similar logic and to delay movements upon trial completion. Only the 50-s testing sequence was retained for analysis.

The stimuli were presented on a gray background with a mean luminance of 61.5 cd/m^2^, equal to the mean luminance of the test images. The monitor was a Display++ LCD with a 120 Hz screen refresh rate, gamma-corrected based on calibrations obtained with a PhotoResearch PR655 spectroradiometer, and controlled by a standard PC. The monitor was viewed at a distance of 80 cm, such that the images subtended a mean width/length of 5.26 degrees of visual angle. To reduce size-specific responses, the size varied randomly over a range from 92-108% of the size of the original image in 4% steps at each image presentation (Dzhelyova & Rossion, 2014). Stimuli were presented over Java SE Version 8. Viewing was binocular and in a room illuminated only by the experimental and acquisition computer displays.

### Task

Participants were instructed to attend to the images presented while fixating on a centrally presented fixation cross, which was present throughout the entire testing sequence, superimposed on the images. To encourage fixation and sustained central attention, the participants’ task was to press on the space bar each time they detected a brief shape change (250 ms) of the cross to an open circle. This occurred 8 times in each trial, at random intervals above a minimum of 500 ms. Participants were naïve to the experimental manipulation.

### EEG acquisition

EEG was acquired with a 128-channel BioSemi ActiveTwo EEG system. This systems uses Ag-AgCl Active-electrodes, with default electrode locations centered around nine standard 10/20 locations on the primary axes, including a reference feedback loop consisting of two additional channels (a common mode sense and driven right leg; BioSemi B.V., Amsterdam, Netherlands; for exact coordinates, see http://www.biosemi.com/headcap.htm). The BioSemi electrodes were relabeled to closely match those of the more conventional 10/5 system (Oostenveld & Praamstra, 2001; for exact relabeling, see Rossion et al., 2015, Figure S2). Additional electrooculogram (EOG) signals were recorded from four flat-type Active-electrodes, positioned above and below the right eye and lateral to the external canthi. Following setup of the EEG system (including insertion of a conductive gel), the offset of each electrode was held below 50 mV. The recordings were saved at a sampling rate of 512 Hz.

### EEG analysis: Preprocessing

Data were (pre)processed with Letswave 6, an open source toolbox (http://nocions.webnode.com/letswave), running over MATLAB R2019b (MathWorks, USA). Data were filtered with a fourth-order, zero-phase Butterworth band-pass filter, with cutoff values below 0.1 Hz and above 100 Hz; to remove contamination from electrical noise, a fast-Fourier transform notch filter was also applied at 60 and 120 Hz, with a width and slope of 0.5 Hz. To correct for muscular artifacts related to eye-blinks, independent-component analysis was applied to remove a single component accounting for blink activity in participants blinking more than 0.2 times/s (6 participants in Experiment 1; overall, M = 0.16 blinks/s; SD = 0.15 blinks/s; 5 participants in Experiment 2; overall, M = 0.11 blinks/s; SD = 0.10 blinks/s). To correct for artifact-contaminated channels (containing deflections beyond ±100 µV in two or more testing sequences), these channels (six or fewer per participant; Experiment 1: M = 3.4 channels; Experiment 2: M = 2.7 channels) were linearly interpolated with 3-5 symmetrically-surrounding neighboring channels. After filtering and artifact correction, data were re-referenced to the common average of the 128 EEG channels.

### EEG analysis: Frequency-domain transform

Preprocessed, individual sequences were isolated in separate 50-s epochs, and averaged in time by condition, to selectively reduce activity not phase-locked to stimulus presentation. These data were transformed into the frequency domain by means of a Fast Fourier Transform (FFT) for amplitude. This spectrum was normalized by the number of samples output; it had a range of 0-256 Hz and a resolution of 0.02 Hz.

### EEG analysis: Harmonic frequencies-of-interest

As mentioned previously, responses to objects (i.e., yellow-selective responses in Experiment 1; and green-selective responses in Experiment 2) are predicted at 1 Hz and its specific harmonics, while general responses to visual object presentation are predicted at 4 Hz and its harmonics. While responses may occur at harmonics beyond the fundamental frequency, they are expected within a limited range, specific to the type of response occurring (e.g., Retter & Rossion, 2016; Jacques, Retter, & Rossion, 2016).

In order to determine the frequency ranges of interest here, the signals were pooled across all EEG channels and grand-averaged across participants. These data were then assessed for significance at all harmonics of the fundamental 1 Hz (color-specific) and 4 Hz (stimulus-presentation) responses for each condition, by means of Z-scores (Z > 1.64; p<.05; calculated at each frequency bin, x, with a local baseline defined by the 20 surrounding frequency bins: Z = (x - baseline mean) / baseline standard deviation; e.g., Srinivasan et al., 1999; Retter & Rossion, 2016). The maximal harmonic frequency range with contiguous significance, exempting one harmonic, in either condition of Experiment 1 was identified and used in subsequent analyses: for the color-specific responses, this range spanned 1 – 25 Hz. Note that harmonics coinciding with the stimulus-presentation responses within this range were excluded. For the stimulus-presentation responses, this range spanned 4 – 56 Hz. These criteria were relatively insensitive to threshold: only one fewer color-specific harmonic would have been selected had a threshold of p<.01 been used. The frequencies-of-interest were used in Experiment 2, after verification that they were reasonably appropriate: green-selective responses were significant until 23 Hz, and until 68 Hz for stimulus-presentation responses.

### EEG analysis: Region-of-interest (ROI) and subregions

An occipito-parietal ROI was defined *post-hoc* from the maximal activation across both conditions of Experiment 1. This ROI consisted of 24 channels, centered medially and extending symmetrically across the left and right cortices. For the yellow-selective responses, averaged across participants it encompassed 21 of the maximal 24 channels for both the grayscale and color conditions; for the corresponding stimulus-presentation responses, it again encompassed 21 of the maximal 24 channels for the grayscale condition, and 22 for the color condition. For more specific response localization, this ROI was further broken into three subregions: the left (10 channels), middle (4 channels), and right (10 channels) (see **Fig. 4**). This ROI was verified for Experiment 2: it captured 21-22 of the maximal 24 green-selective channels, and 20-21 of the maximal stimulus-presentation channels, in each condition.

### EEG analysis: Quantification and statistics

Responses were examined across all the EEG channels and harmonic frequencies of interest; however, to summarize the results in quantification and statistical analyses, the primary analyses focused on data collapsed across the ROI (and subregions), through channel-averaging, and frequencies-of-interest, through amplitude summation (Retter & Rossion, 2016). Data were grand-averaged across participants for description and display, as well as for a group-level tests of response significance on the summed harmonic responses (Z-scores: Z > 2.32; p<.01). Note that after harmonic responses were combined, a local baseline-correction was applied (given the variable noise level across the frequency spectrum), in the form of a baseline-subtraction, with a baseline defined by the 20 surrounding frequency bins, after excluding the local minimum and maximum (e.g., Rossion et al., 2012; Retter & Rossion, 2016).

To statistically compare the responses across conditions at the occipito-parietal ROI, in Experiment 1, one-tailed, paired-sample t-tests were performed, with the prediction that larger amplitude responses would be produced in the color than grayscale condition. A one-way ANOVA was performed to compare the three conditions of Experiments 2 and 3, followed up by one-tailed paired-sample t-tests comparing the color and grayscale condition of Experiment 2 (as in Experiment 1), as well as the color and incongruent conditions. To compare the spatial distribution of responses across the scalp, a repeated-measures analysis of variance (ANOVA) was performed, with factors of *Condition* (two levels in Experiment 1: grayscale and color; three levels in Experiments 2 and 3: grayscale, color, and incongruent) and *Subregion* (three levels: left, medial, and right). In the case that Mauchly’s test of sphericity was violated, a Greenhouse-Geisser correction was applied.

### EEG analysis: Time domain

Data were analyzed in parallel in the time domain, following preprocessing (e.g., as in Rossion et al., 2012; Retter & Rossion, 2016; Jacques, Retter, & Rossion, 2016). To this extent, the data were first low-pass filtered with a fourth-order Butterworth filter at 30 Hz. To isolate the object responses, the stimulus-presentation responses were selectively removed through a frequency-domain notch filter, applied at all the harmonics of 4 Hz below 30 Hz, with a width and slope of 0.02 Hz. The data were cropped in separate segments for each object presentation, from 250 ms prior- and 750 ms post-stimulus onset. The cropped data segments were baseline-corrected by subtracting the average amplitude in the 250 ms preceding stimulus onset, a time window corresponding to one stimulus-presentation cycle. To avoid contamination from eye movements, data segments containing deflections of ± 125 µV in any EOG channel were rejected. Data segments were then averaged by condition. To determine when color-specific response deflections significantly differed from baseline (0 µV), independent t-tests against zero were performed at every time bin from stimulus onset to 750 ms post-stimulus onset; to reduce the chance of false-positive due to the high number of comparisons, a conservative threshold of p<.01 was selected, and consecutively criteria of 15 ms (9 consecutive sampling bins) was applied. Similarly, paired-sampled t-tests were applied to determine when the responses from the color and grayscale conditions differed from each other, as well as the color and incongruent conditions in Experiment 2. For description and display, data were grand averaged across participants.

## Results

### Frequency domain: amplitude spectra

Responses to visual stimulus presentation were evident as high-amplitude peaks in the frequency domain at 4 Hz and its harmonics. More importantly, color-specific responses were also evident as peaks at 1 Hz and its specific harmonics, for every condition (**Fig. 2a** for an example: see **Fig. S1** for all conditions). To describe these complete responses, the harmonics were summed and baseline-subtracted, from 1 to 25 Hz for color-specific responses, and from 4 to 56 Hz for stimulus-presentation responses (Experiment 1: **Fig. 2b**; Experiment 2: **Fig. 2c**; see Methods).

**Figure 2.**
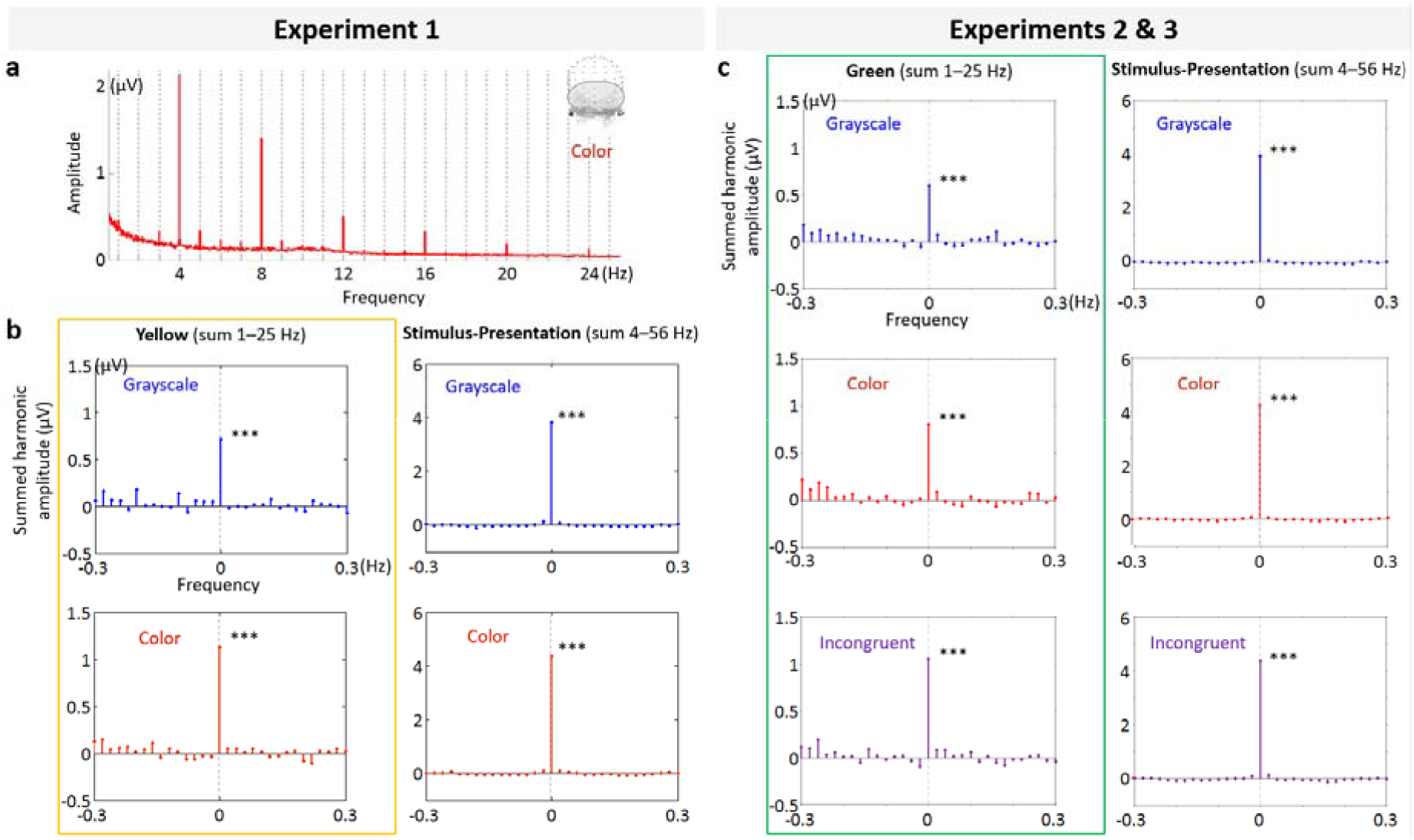
**a)** The frequency-domain amplitude spectra for the color condition of Experiment 1, plotted over the occipito-parietal ROI. Dotted vertical lines indicate the frequencies of 1 Hz and its harmonics. (For corresponding spectra of all conditions, see **Fig. S1**.) **b & c)** Summed baseline-subtracted harmonic responses: the frequency-of-interest and its harmonics are combined and plotted relative to the stimulation frequency (0 Hz corresponds to 1 Hz and its specific harmonics (left) or 4 Hz and its harmonics (right), with the surrounding frequency bins included for comparison); noise level is at 0 µV. **Key**: *** = Z>3.1, p<.001.

### Frequency domain: amplitude quantification

Critically, the color-specific responses to objects at 1 Hz and its harmonics were robust even when the color-associated objects were shown in grayscale. The amplitude of the color-specific summed-harmonic response to objects in grayscale was about 70% of that for objects in color (Experiment 1: 63%; Experiment 2: 75%) over the occipito-parietal ROI (**Fig. 3a**; **Table S1a**). In comparison, for stimulus-presentation, the grayscale response was about 90% that of color (Experiment 1: 81%; Experiment 2: 93%; **Fig. 3b**; **Table S1b**).

**Figure 3.**
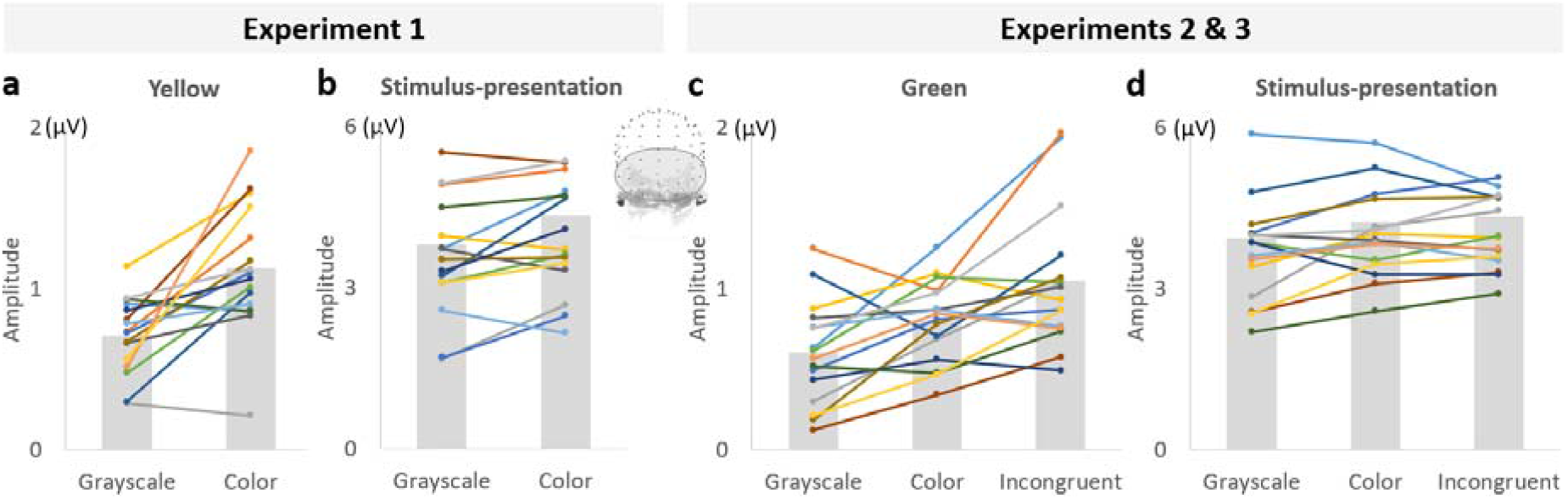
Summed-harmonic, baseline-subtracted response amplitude (µV) over the occipito-parietal ROI, at the group- (gray bars) and individual- (colored lines) levels. **a)** Yellow-selective responses (1-25 Hz) from Experiment 1. **b)** Stimulus-presentation responses (4-56 Hz) from Experiment 1. **c)** Green-selective responses (1-25 Hz) from Experiments 2 and 3. **b)** Stimulus-presentation responses (4-56 Hz) from Experiments 2 and 3.

This amplitude difference between the yellow-selective responses in the color and grayscale conditions was statistically significant in both experiments: Experiment 1, t_15_ = 4.22, *d* = 1.29, *r* = 0.54, *p* = .0004; Experiment 2, t_15_ = 2.94, *d* = 0.70, *r* = 0.33, *p* = .0051 (following a significant one-way ANOVA, F_2,45_ = 6.94, *η*_*p*_^*2*^ = 0.24, *p* = .0024). At the individual level, significant color-specific responses at the occipito-parietal ROI were found in all but one participant in both conditions in Experiment 1, and in all but three participants in the grayscale condition in Experiment 2, underlying the reliability of the main finding that both grayscale and color images are sufficient to elicit color-specific responses to color-associated objects. With regard to the corresponding stimulus-presentation responses, these were significantly different in Experiment 1, t_15_ = 1.75, *d* = 1=0.32, *r* = 0.16, *p* = .012, but a one-way ANOVA in Experiments 2 and 3 indicated no significant differences across conditions, F_2,45_ = 0.41, *η*_*p*_^*2*^ = 0.018, *p* = .66.

In the other direction, the color-specific response to incongruently colored objects in Experiment 3 was 131% of that to correctly-colored objects, t_15_ = -3.10, *d* = -0.70, *r* = -0.33, *p* = .0037. However, there was no appreciable difference in the corresponding (color vs. incongruent) stimulus-presentation responses (**Fig. 3c&d**; **Table S1**).

### Frequency domain: topography

For a more detailed spatial investigation of these responses, the scalp topographies were plotted (**Fig. 4a&c**) and the single occipito-parietal ROI was decomposed into its left, right, and medial subregions (**Fig. 4b&d**). Across the first two experiments, the scalp topographies showed more lateralized color-specific responses for grayscale than color conditions (**Fig. 4a&c**). Although responses appeared right lateralized for yellow-associated object responses, and left lateralized for green-associated object responses overall, there was great inter-individual variability: indeed, in Experiment 2, there were not more left-than right-lateralized participants (**Fig. S2**).

**Figure 4.**
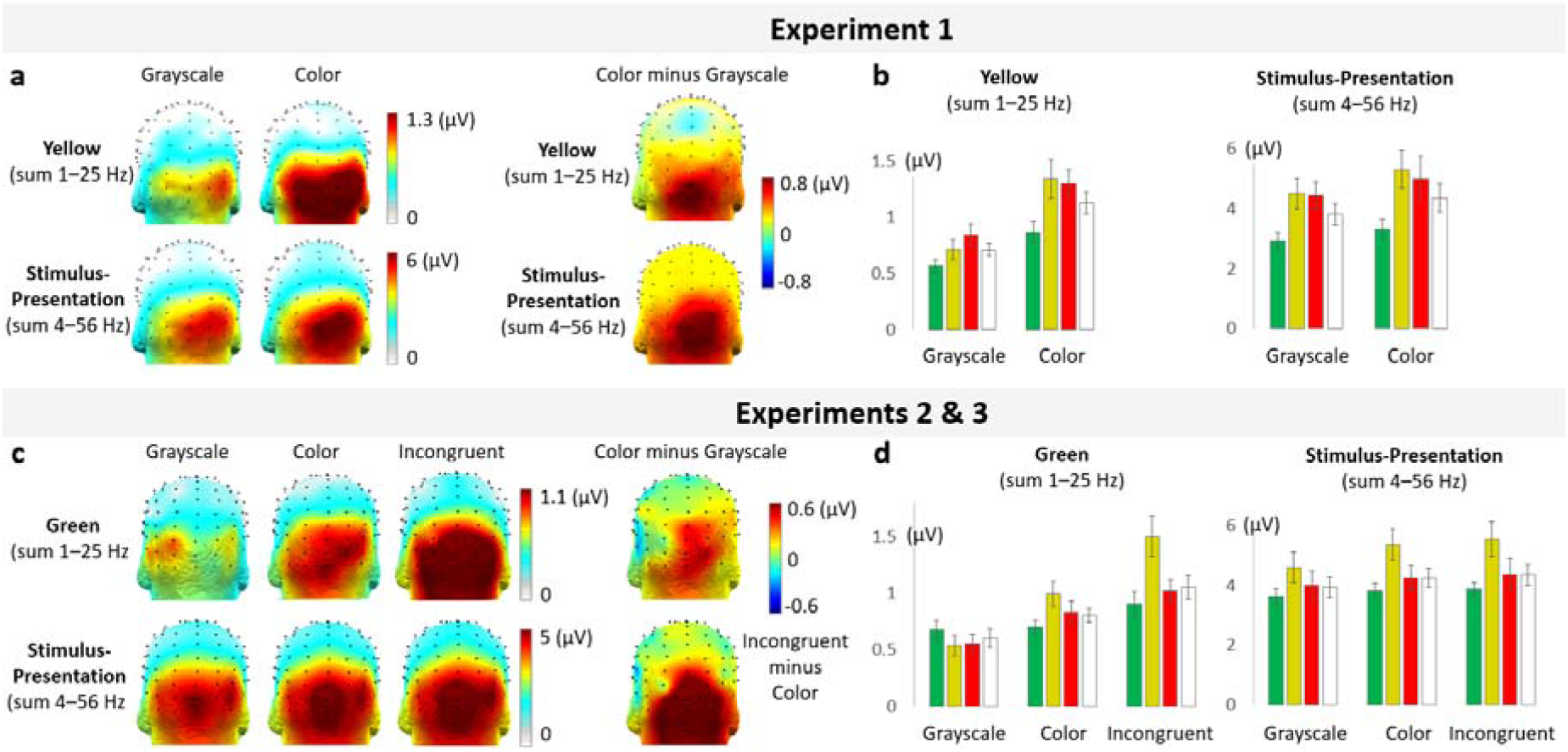
**a & c)** Summed-harmonic response scalp topographies in the grayscale and color conditions (Experiments 1 and 2), and incongruent condition (Experiment 3), as well as their differences. **b & d)** Mean summed-harmonic responses at the occipito-parietal ROI and its three subregions: left, medial, and right.

In the subregions of interest, in the middle the response in the grayscale condition was about 54% of that of the color condition across the first two experiments, while over the combined left and right subregions, it was about 74% (middle: 53%-54% across experiments; left and right: 82-65% across experiments). In comparison, the stimulus-presentation responses were more similar in the grayscale relative to color conditions, and less specifically over the middle subregion: about 85% middle, and 92% over the left and right subregions (middle: 84%-56% across experiments; left and right: 95-88% across experiments).

In Experiment 1, these spatial differences for color-specific responses across conditions were confirmed in a repeated-measures ANOVA, with factors of *condition* and *subregion*, which yielded a moderate interaction between these factors, *F*_2,30_ *=* 4.03, *p* = .042, *η*_*p*_^*2*^ = 0.37. Note that the main effect of this ANOVA replicated the large difference of *condition, F*_1,15_ *=* 18.7, *p* = .001, *η*_*p*_^*2*^ = 0.56; the main effect of *subregion* was not robust, *F*_1,30_ = 2.96, *p* = .084, *η*_*p*_^*2*^ = 0.30. Similarly, in Experiments 2 and 3, there was an interaction between *condition* and *subregion, F*_4,15_ *=* 6.33, *p* = .0003, *η*_*p*_^*2*^ = 0.30, and a substantial main effect of *condition, F*_2,15_ *=* 18.6, *p* < .0001, *η*_*p*_^*2*^ = 0.55. There was also a main effect of *subregion, F*_2,15_ *=* 6.63, *p* = .0041, *η*_*p*_^*2*^ = 0.31.

### Temporal dynamics

To investigate the temporal dynamics of the color-specific responses, the data were analyzed in the time domain, in terms of event-averaged responses time-locked to the onset of yellow-associated (Experiment 1), green-associated (Experiment 2), or incongruent green (Experiment 3) object stimulus presentation, over the occipito-parietal subregions (see Methods).

Initially, the responses to yellow- or green-associated/incongruent objects were obscured by the responses to other objects (appearing every 250 ms, i.e., at – 250 ms, 250 ms, and 500 ms here; Experiment 1: **Fig. 5a;** Experiments 2 and 3: **Fig. S3a**). To isolate color-specific object responses, the waveforms were notch-filtered at the stimulus-presentation rate, 4 Hz, and its harmonics. Note that the response aspects common to the stimulus presentation are removed by the notch filter, such that if no differential response to the target stimuli at 1 Hz is recorded, no substantial deflections from the baseline would be present.

**Figure 5.**
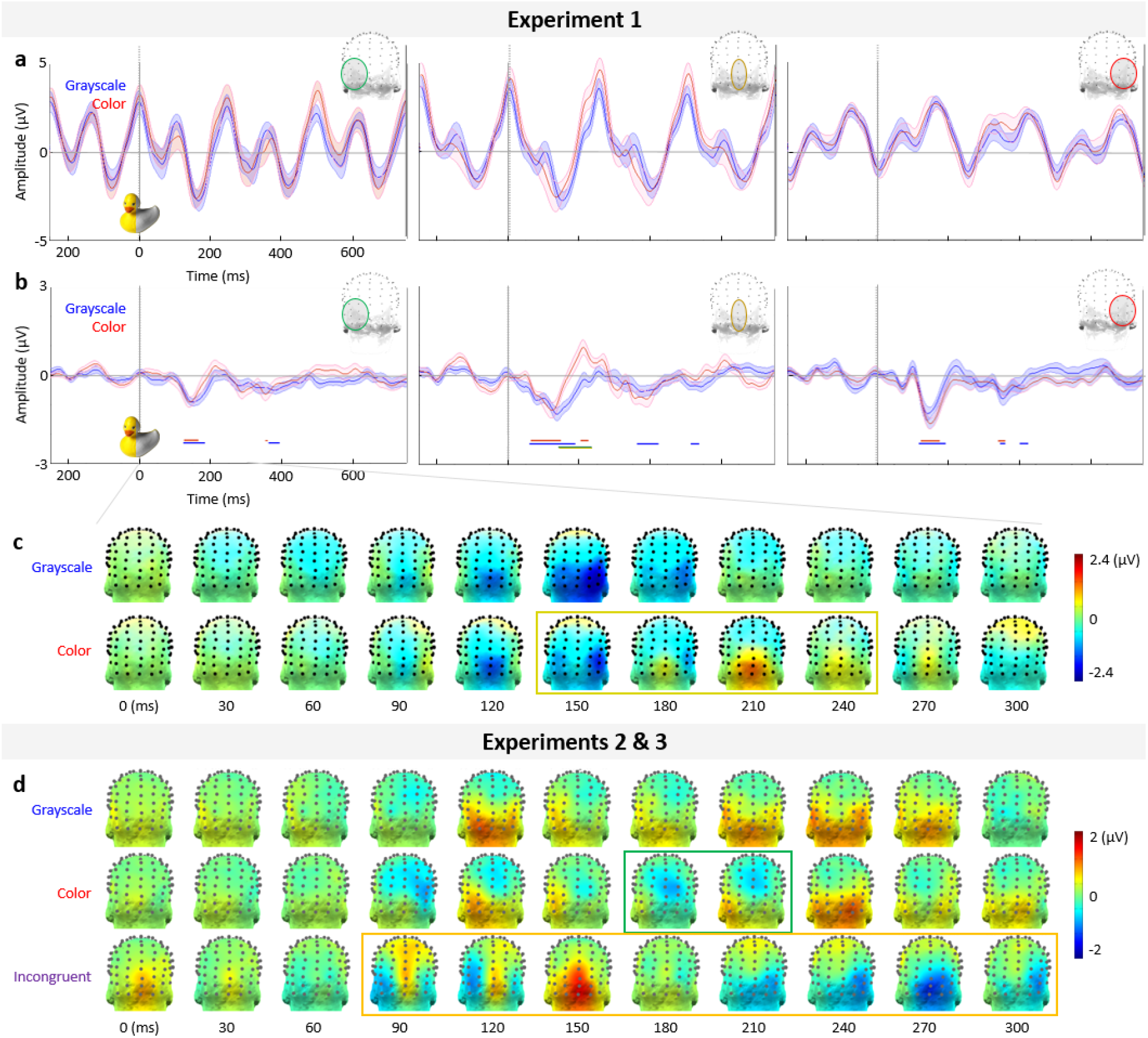
Temporal dynamics. **a)** Time-domain responses in the grayscale (blue waveform) and color (red waveform) conditions, related to yellow-associated object stimulus onset (0 ms; exemplified with a yellow/gray rubber duck; in Experiment 1), at the three scalp subregions (left, middle, and right panels, respectively). These responses reflect the responses to all objects (presented at 4 Hz, i.e., every 250 ms) as well as the responses specific to yellow-associated objects. The dark waveforms are the average across subjects, with the shaded areas indicating ± 1 SE. **b)** After filtering out the general visual EEG responses at 4 Hz and its harmonics, the time-course of the yellow-associated object responses may be isolated. This panel is plotted as in (a), except with a different amplitude scale. Additionally, the time windows of significant deflections for each condition are indicated by solid lines below the waveforms in the corresponding color by condition; significant differences across conditions, occurring only at the middle subregion, are indicated in yellow-green. **c)** Response scalp topographies across time: sampled every 30 ms from stimulus onset. The approximate time window of significant differences between conditions, i.e., a positive deflection over the middle subregion in the color condition, is highlighted with a yellow outline over that condition. **d)** Response scalp topographies across time, for Experiments 2 and 3, plotted as in (c), with a green outline over the color condition; differences between the color and incongruent conditions are outlined in orange over the incongruent condition.

There were significant deflections in both the grayscale and color conditions, in a range of about 60 to 670 ms post-stimulus onset across the first two experiments (**Fig. 5b; Fig. S3b**). In Experiment 1, these responses first onset in a negative deflection over the middle subregion (grayscale: 61 ms; color: 65 ms). A negative deflection was also seen later over the left and right subregions, first reaching significance at 119 and 125 ms over the right subregion in the grayscale and color conditions, respectively, and at 123 and 127 ms over the left subregion. In total, there were no differences in the onset latency of the yellow-associated response across the grayscale and color conditions. In Experiment 2, color-specific responses first onset in a negative deflection over the middle subregion in the color condition (172 ms), but in a positive deflection over the left subregion in the grayscale condition (109 ms). Note that the differences in the temporal dynamics of the responses specific to yellow and green objects (in Experiments 1 and 2, respectively) supports the presence of color-specific responses in the frequency domain.

Importantly, the significant differences between the color and grayscale conditions occurred in a later time window, from 140-230 ms post-stimulus onset, over the middle subregion (Experiment 1: 141-234 ms; Experiment 2: 174-211 ms; paired sample t-tests; p<.01; **Fig. 5b; Fig. S3**). In both experiments, this was present as a deflection present in the color but not grayscale condition: Experiment 1, a positive deflection peaking at 211 ms; Experiment 2: a negative deflection peaking at 188 ms (**Fig. 5c&d**). There were no other appreciable differences across conditions in either the subregion waveforms, or the scalp topographies, in either experiment.

In Experiment 3, there were also significant, early, specific deflections to incongruently-colored green objects. However, these presented first over the left subregion, at 68 ms, shortly followed by the middle subregion, at 76 ms (**Fig. S3b**). The responses to incongruently-colored green stimuli also showed relatively early, significant differences from those to congruently-colored green stimuli, first emerging at 98 ms over the left subregion, followed by 125 ms over the middle subregion (**Fig. 5d; S3b**).

## Discussion

### Summary

We designed an experiment to isolate selective responses to objects of one color category (yellow in Experiment 1; green in Experiments 2 and 3), presented at a rate of 1 Hz within a 4 Hz stream of otherwise non-periodic object colors. We found color-specific responses reflecting the differing spatiotemporal dynamics of EEG responses to different colors (e.g., Regan, 1966; Riggs & Sternheim, 1968; Allison et al., 1993; Anllo-Vento, Luck & Hillyard, 1998), for stimuli with strongly associated colors shown in actual color. Critically, these automatic, color-specific responses were also generated from grayscale stimulation through object knowledge alone (Experiments 1 & 2), and were modulated for incongruently-colored objects (Experiment 3). The early onset of these color-specific responses in all stimulus conditions contrasted with a later role of actual color (after 140 ms post-stimulus onset). These results support distinct roles of color at early and late processing stages: they are in line with a framework in which object color knowledge is part of the early visual shape representation, while suggesting that actual color plays a later, specific role in visual processing.

### Color-specific EEG responses to actually colored stimuli

Selective neural responses were recorded at 1 Hz and its harmonics to yellow (Experiment 1) or green (Experiment 2) objects shown in their associated color. This demonstrates a neural response to the consistent presentation frequency of the target color that is distinct from the responses to non-periodic presentations of three other object color categories, also shown in their associated color (color condition; **Fig. 2**). Different responses to different colors have been reported in a few previous EEG studies (e.g., Regan, 1966; Riggs & Sternheim, 1968; Anllo-Vento, Luck & Hillyard, 1998; Retter et al., 2020). This is unsurprising given differences in amplitude and latency as a function of cone-opponent cortical inputs (e.g., Robson & Kulikowski, 1998; Rabbin et al., 1994; Lee et al., 2009). However, these results have been previously reported only at the group level, typically with a small number of electrodes, and were largely descriptive: here, we objectively quantify color-specific responses in the frequency domain which are significant at the individual participant level (in 15/16 participants in Experiment 1; and 16/16 participants in Experiment 2; see **Fig. 3a&c**). In our results, the high sensitivity to the target color is likely enabled by the extensive spatial coverage of high-density EEG (given the variability in the response scalp topography across individual participants), as well as the high signal-to-noise ratio afforded by frequency tagging (Regan, 1966; Norcia et al., 2015).

There are several reasons to conclude that this color-specific response does not emerge as an artifact of our paradigm. First, there was no confound of frequency of occurrence across object color categories: all object images appeared equally as often in each sequence on average. While all object colors thus on average were occurring at a rate of 1 Hz, we empirically demonstrated that no response would occur at 1 Hz and its harmonics without a specific periodically-presented color: in a control condition added with the last five participants of Experiment 1, we presented all the colored objects non-periodically, and found no 1-Hz response (**Fig. S4**). Secondly, the “oddball-like” paradigm used here has been demonstrated to be immune to temporal predictability of the target stimuli (Quek & Rossion, 2017), and to show discriminable responses to different object image types (Jacques, Retter & Rossion, 2016). Third, the yellow-selective response we record in this paradigm is similar in terms of scalp topography to a yellow/gray asymmetry response reported in a previous study (Retter et al., 2020). Finally, a specific response was present consistently for both yellow and green objects, and was modulated by incongruent color in Experiment 3, again suggesting that the responses at 1 Hz and its harmonics do appear to capture color-specific responses.

That the amplitude of the color-specific response to colored objects was larger than that to grayscale objects may represent the contribution of the actual color response amplitude, in line with neural responses to non-object color stimuli. It should be remembered that the color-specific responses recorded here are *differential* responses to objects presented at 1 Hz, *vs*. object presentation in general, at 4 Hz. The color-specific responses are thus more reduced for grayscale vs. color stimuli than would be expected from a general effect of color, as indexed by the responses at the stimulus-presentation rate (again, the grayscale responses were about 70% of the amplitude of actual color responses for the color-specific responses over the occipito-parietal ROI, but about 90% for the stimulus presentation responses; **Fig. 3**; see also Or, Retter & Rossion, 2019, for no general effect of color at 12 Hz). On the other hand, when stimuli were shown in incongruent colors in Experiment 3, the largest response was produced, (**Fig. 4d;** as in Lu et al., 2010; Lloyd-Jones et al., 2012; but not in Provebio et al., 2004), possibly indicating a conflict of the associated and actual colors.

### Color-associated objects elicit color-specific EEG responses to grayscale images

We recorded selective responses to yellow- (Experiment 1) and green- (Experiment 2) associated objects at 1 Hz and its harmonics, even when all the objects were shown in grayscale (grayscale condition; **Fig. 2**). As addressed in the Introduction, previous studies have shown that color memory influences visual object perception, through a convergence of various experimental approaches. Indeed, color-specific responses to grayscale stimuli have been previously reported with functional magnetic resonance imaging (Bannert & Bartels, 2013; Vandenbroucke et al., 2016) and MEG (Teichmann et al., 2018; 2019). Here, the report of color-specific responses to grayscale objects with EEG, novel to our knowledge, supports and extends these previous studies.

Previous EEG studies have instead measured effects of color memory that are non-specific to a color category, i.e., contrasts of color *vs*. grayscale or color *vs*. incongruent color objects, averaging across color categories (8 different color categories: Proverbio et al., 2004; color category uncontrolled among 96 objects: Lu et al., 2010; color category uncontrolled among 54 high color-diagnostic objects: Bramao et al., 2012b; color category uncontrolled among 150 objects: Lloyd-Jones et al., 2012). Not only did these previous EEG studies report inconsistent differences in the amplitude of color *vs*. grayscale objects responses, but these effects cannot be taken as reflecting color-specific responses associated with objects. In fact, when only color *vs*. grayscale object responses are compared, the response differences could be more related to a general effect of color, as we also observe on the 4 Hz stimulus presentation rate in our study (i.e., a grayscale response of about 90% that of color). Additionally, no previous study quantified the difference in response amplitude between implied *vs*. actual color.

The presence of color-specific responses to grayscale objects here, about 70% of the amplitude of response to color objects, suggests that color-specific responses may arise from responses to the objects, of which associated color is one common aspect. That is, shape alone may automatically trigger full neural representations of objects, including their associated color knowledge. For example, the brain’s response to seeing the shape of a yellow rubber duck may automatically activate a network of cortical regions representing this object through its attributes, including its yellow color. In this vein, it has been suggested that color memory responses originate from color-selective regions evoked from object shape associations (e.g., Slotnick, 2009, recording learned color-shape associations of abstract visual shapes; see also Simmons et al., 2007). Another possibility is that color knowledge responses originate from regions close to but external from color-selective regions, which may have been influenced by color associations (e.g., Martin et al., 1995, recording shape associations to the presentation of color words; note that similar neural data is interpreted differently by Slotnick, 2009). Thus, it remains unknown whether these color memory responses originate from color-selective cortical regions, or whether they are object-selective responses that have been selectively shaped by learned color associations.

Importantly, this reduced yet still substantial response amplitude should not be interpreted as the perceptual experience of color in grayscale images. Indeed, under most conditions, grayscale images of color-associated objects actually do appear gray (with the exception of small effects of memory color biases, produced in an unnatural context: e.g., Hansen et al., 2006, reporting 4-13% relative scaling; Lee & Mather, 2019, reporting weak chromatic adaptation effects to achromatic implied color stimuli). However, the presence of color-specific brain responses does not imply that one is actively perceiving yellow when looking at a grayscale rubber duck. Similarly, large amplitude brain responses to perceiving eyes alone does not imply one is perceiving a whole face (Bentin et al., 1996), and the pattern and rate of “mirror neuron” activity for visually observing motion does not imply one perceives themselves moving (e.g., Kilmer & Lemon, 2013). Instead, again, our results suggest that perceiving information about an object may automatically activate representations of other attributes associated with the object, even if they are absent from the current stimulus.

### Equal onset-latencies are followed by a late (∼140-235 ms) actual color difference

The stage at which color associations may arise from object knowledge remains highly controversial. In the time-domain analyses, we show that color-specific responses to color-associated grayscale objects emerge at the same time as those to actually colored objects (**Fig. 5**). This suggests that neural responses to objects, derived from spatial cues alone, are rapidly and automatically influenced by their remembered color associations: objects are categorized by color with no delay between grayscale and colored images. These findings support an early role of color knowledge in object recognition, at least for familiar objects routinely associated with specific colors (i.e., “color-diagnostic” objects).

In contrast, the responses to actually colored objects diverged from those to grayscale objects only later, in the form of an additional deflection present only in the actual color condition (**Fig. 5b-d; Fig. S3b**). This deflection was present within about 140-235 ms post-stimulus onset; its peak was on average at approximately 200 ms, across experiments. Spatially, this deflection was centered over the middle occipital channels, in agreement with the middle subregion producing the largest amplitude advantage for the color condition in the frequency-domain analysis (e.g., in Experiment 1, compare the scalp topography of the differences across conditions in **Fig. 4a** with that of this time-domain component in **Fig. 5c**). The latency of these deflections is in line with the earliest chromatic visually evoked potentials to isoluminant stimuli reported with EEG in the human brain (from about 100-120 ms: e.g., Rabin et al., 1994; Gerth et al., 2003: the C_II_-C_III_ components; Nunez, Shapley & Gordon, 2017). Thus, it is possible that these late deflections are related to typical chromatic processing.

Note that despite this relatively late neural signature specific to chromatic processing, color may still play an earlier role in combination with luminance (e.g., see the C_I_ component in Gerth et al., 2003). One possibility is that such a color-luminance effect may not be strongly color-specific (at the population level), that is to say, not substantially aiding in differentiating actually colored yellow, green, blue, and red objects here, and thus not contributing to the 1 Hz responses in our design. In that case, this effect may have had an impact more generally at 4 Hz and its harmonics (although no latency difference was observed (**Fig. 5a);** however, the 4-Hz response amplitude was greater for color than grayscale images, **Fig. 3b**; **Table S1b)**. Another, not mutually exclusive possibility, is this early effect of color had only a weak impact here, since our stimuli could be rapidly processed in any case (they were centrally presented, cropped images, without shadows or occlusion, 250 ms stimulus-onset asynchronies, etc.; e.g., see Allen, 1879; Walls, 1942, p. 463; Elsner, 1978; Frome, Buck, & Boynton, 1979; De Valois & Switkes, 1983; Gegenfurtner & Rieger, 2000; Bramao et al., 2012). Finally, moderately late differences to incongruently-colored and congruently-colored green stimuli were found: over the middle occipito-parietal subregion, these first occurred at 125 ms (**Fig 5d; Fig S3)**. The differences in responses to incongruent vs. color objects may reflect the interference of color-shape integration in object processing, which again may be bound in this time window in visual processing (see also Lu et al., 2010; Lloyd-Jones et al., 2012; Moutoussis, 2015).

Together, these findings relate to the debate as to which stage color has an effect on object recognition, in combination with object shape, as addressed in the Introduction. On one hand, we provide evidence that color associations are integrated into objects’ (shape-driven) representations or associations early on. This finding is in line with some earlier EEG studies (e.g., Lu et al., 2010; Bramao et al., 2012b). It is also in line with behavioral studies showing a color advantage in recognizing objects or scenes with associated colors, even at the shortest image presentation times (from 10 to 1000 ms per item: Bruner & Postman, 1949; from 16-64 ms per image: Gegenfurtner & Rieger, 2000; from 13 to 80 ms per image: Hagmann & Potter, 2016), providing indirect evidence that color has an early impact on object processing. Further indirect evidence has been provided by behavioral studies that show a consistent color advantage in accuracy of object recognition across trials with relatively early or late response times (Rossion & Pourtois, 2004; Hagen et al., 2014). There is evidence for an early integration of color with shape in visual processing (e.g., Thorell, De Valois & Albrecht, 1984; Johnson, Hawken & Shapley, 2001; Seymour et al., 2010), which may contribute to the rapidity of these learned associations, and enable the modulation of early (and later) color-specific responses in the case of incongruently colored objects.

On the other hand, the present results suggest that actual color does have a secondary role to shape in visual object recognition (e.g., Grossberg and Mingolla, 1985; Biedermann 1987). This is also in line with previous studies, e.g., in color having a late effect on enhancing image representation in memory (Gegenfurtner & Rieger, 2000; for other late effects of color on object categorization: Provebio et al., 2004; Yao & Einhauser, 2008; Bramao et al., 2012b; Lloyd-Jones et al., 2012; Or, Retter & Rossion, 2019). It is also in line with the separation of color and shape in visual processing (e.g., Treisman, 1982; Cavanagh, Tyler & Favreau, 1984; Zeki & Shipp, 1988; Livingstone & Hubel, 1984; Pinna, Brelstaff & Spillmann, 2001). By distinguishing color *knowledge* from *actual* color (similar to color *identification* from color *contrast perception*: Johnson & Mullen, 2016; or *stored color knowledge* from *surface color*: Joseph & Proffitt, 1996), it may be possible to distinguish two distinct roles of color in visual object recognition. Perhaps contrary to intuition, the present results suggest that color knowledge occurs at early stages of visual processing, even preceding signatures of actual color(-specific) perception.

Our findings are not compatible with hierarchical views of color first being processing selectively in early visual areas, and only later being influenced by “top-down” feedback from color memory expectations at later stages (e.g., Tanaka, Weiskopf, & Williams, 2001; Hansen et al., 2006; Hanslmayr et al., 2008; Bramao et al., 2012). Instead, our findings may be interpreted in a framework in which the visual world is perceived initially through associative recognition: i.e., the details of the visual world, captured through sensory processing, are not fully processed in advance of stimulus recognition, which incorporates associated aspects of an object in the human brain (see Helmholtz, 1867; Gregory, 1966; Sergent, 1986; Thompson, 1995; Lotto & Purves, 2002). Note that such a model was also proposed by Joseph & Proffitt (1996: Model B, Fig. 5), in the context of color knowledge having a larger effect than actual color in a behavioral object recognition task, although conservatively stated in terms of influence rather than actual processing time. Here, electrophysiological data support that object color knowledge may indeed affect our earliest perception of color.

## Acknowledgements

This work was supported by grants EY-010834 and P20 GM103650 to MW, and grant FC7159 to TLR. Thanks to those who gave feedback on earlier versions of this work: Alex Wade, Olivier Collignon, Valérie Goffaux, and Dennis Mathew.

## Supplementary Material

**Supplemental Figure 1.**
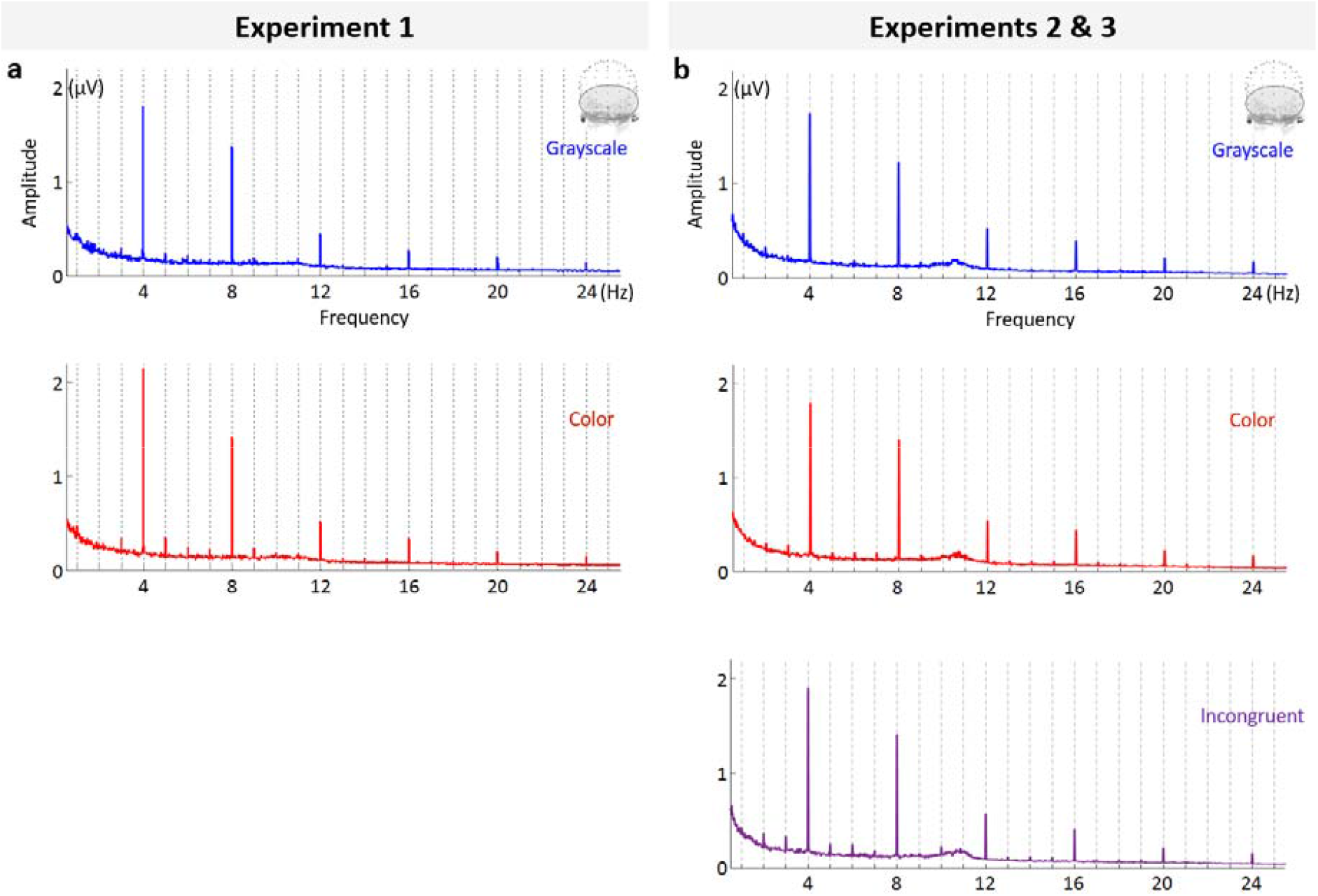
Frequency-domain amplitude spectra for all the of the experimental conditions, over the occipito-parietal ROI. **A)** Experiment 1. **B)** Experiments 2 & 3.

**Supplemental Figure 2.**
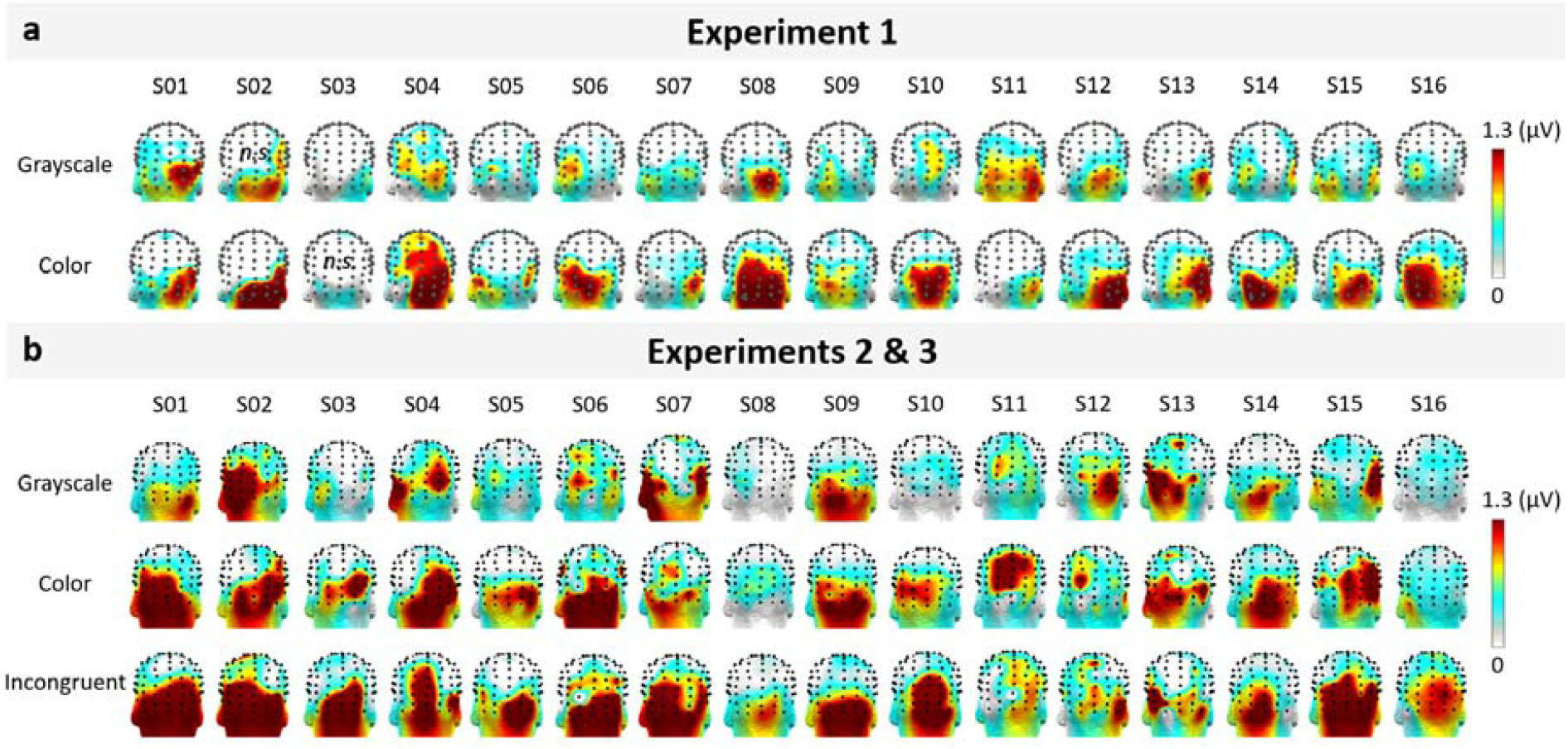
Individual participants’ color-specific summed-harmonic response amplitudes (1-25 Hz) at the occipito-parietal ROI. **a)** Experiment 1. **B)** Experiments 2 & 3. Note that the participants are different across Experiments 1 and 2.

**Supplemental Figure 3.**
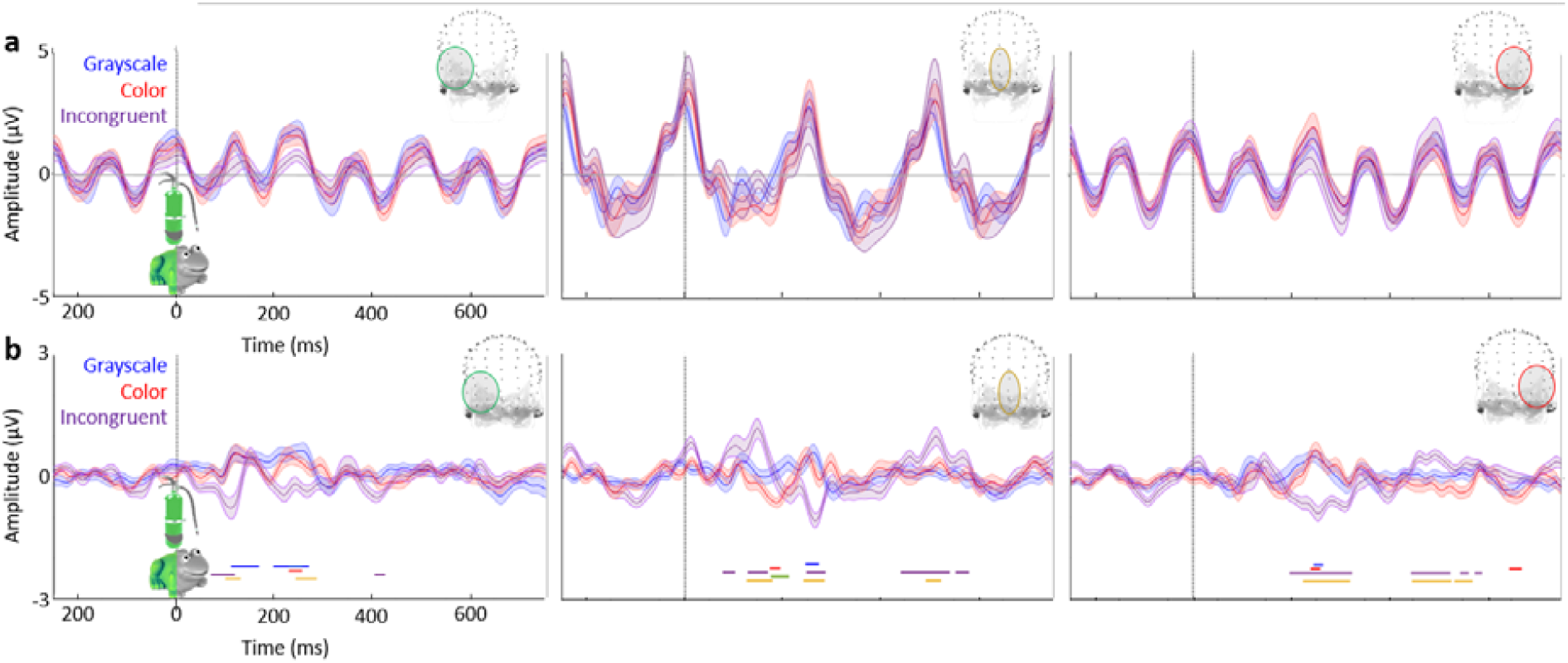
Time-domain responses in Experiments 2 and 3 (plotted as in **Fig. 5**). **a)** Responses including those to stimulus-presentation at 4 Hz (every 250 ms), as well as those specific to green-associated (Experiment 2) or incongruently-green (Experiment 3; purple waveform) objects at 1 Hz. **b)** Responses at 1 Hz, i.e., following a notch-filtering of the stimulus-presentation responses, at 4 Hz and its harmonics.

**Supplemental Figure 4.**
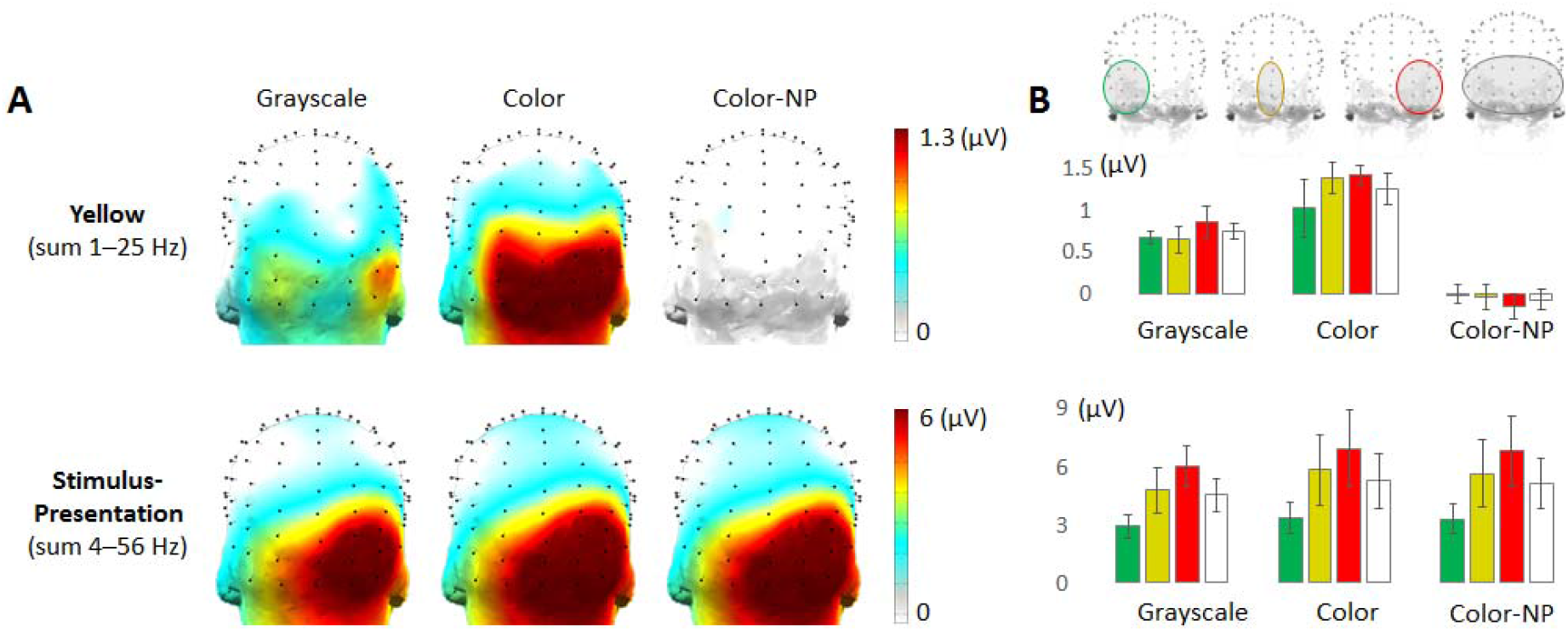
Responses to a color non-periodic (NP) control condition, in which yellow repeated non-periodically throughout the stimulation sequences (N = 5). **a)** No response is present at 1 Hz and its harmonics when no color is presented periodically (“frequency-tagged”) at this rate. **b)** When quantified at the occipito-parietal ROI and its subregions, there is again no response for the non-periodic control condition at 1 Hz and its harmonics, however, note that the stimulus-presentation response in this condition is equivalent to that of the standard periodic color condition.

**Supplemental Table 1.** Summed-harmonic, baseline-subtracted response amplitude (µV) over the occipito-parietal ROI. In parentheses is one standard error of the mean.

|  | a) Color-specific |  | b) Stimulus-presentation |  |
| --- | --- | --- | --- | --- |
|  | Experiment 1 | Experiments 2 & 3 | Experiment 1 | Experiments 2 & 3 |
| Grayscale | 0.71 (0.059) | 0.60 (0.079) | 3.81 (0.360) | 3.94 (0.338) |
| Color | 1.13 (0.098) | 0.80 (0.063) | 4.35 (0.481) | 4.25 (0.306) |
| Incongruent |  | 1.05 (0.11) |  | 4.36 (0.355) |

### Image statistics: color category comparisons

For spatial frequency content, five levels of spectral energy (10, 30, 50, 70, and 90%) were compared over frequencies across color category groups, with no differences produced at any level (all F’s <1.95; p’s > .15; one-way ANOVAs). The amount of energy at high spatial frequencies (defined as over 15 cycles/image) also did not differ across groups, F_3,20_ = 1.36, p = .29, *η*_*p*_^*2*^ = 0.13.

Global contrast factor reflects a contrast measure across a number of resolution levels that is thought to relate to perceptual contrast: this factor did not vary across images (GCF range across color category means: 12.4-14.1, F_3,20_ = 1.03, p = .40, *η*_*p*_^*2*^ = 0.17; one-way ANOVA).

Gist relates to visual spatial forms that can quickly and automatically be extracted from an image, and may be measured with spatial envelopes: differences in gist did not differ across image categories (D range: 0.94-1.06), F_3,68_ = 1.98, p = .13, *η*_*p*_^*2*^ = 0.08 (one-way ANOVA including all paired cross-category differences for each image).

Although images were sized to a common rectangular area, further, the non-background area did not differ across color categories: F_3,20_ = 2.30, p = .11, *η p*^*2*^ = 0.26; one-way ANOVA).

## References

Allen, G. (1879). The colour-sense: its origin and development. Mind, 4(15), 415–421.

Allison, T., Begleiter, A., McCarthy, G., Roessler, E. Nobre, A. C. & Spencer, D. D. (1993). Electrophsyiological studies of color processing in human visual cortex. Electroencephalography and Clinical Neurophysiology, 88(5), 343-355.

Anllo-Vento, L., Luck, S. J. & Hillyard, S. A. (1998). Spatio-temporal dynamics of attention to color: evidence from human electrophysiology. Human Brain Mapping, 6(4), 216–238.

Bae, G. Y., Olkkonen, M., Allred, S. R., & Flombaum, J. I. (2015). Why some colors appear more memorable than others: A model combining categories and particulars in color working memory. Journal of Experimental Psychology: General, 144(4), 744–763.

Bannert, M. M. & Bartels, A. (2013). Decoding the yellow of a gray banana. Current Biology, 23, 2268–2272.

Bartleson, C. J. (1960). Memory colors of familiar objects. Journal of the Optical Society of America, 50(1), 73–77.

Bentin, S., Allison, T., Puce, A., Perez, E., & McCarthy, G. (1996). Electrophysiological studies of face perception in humans. Journal of Cognitive Neuroscience, 8(6), 551–565.

Biedermann, I. (1987). Recognition-by-components: a theory of human image understanding. Psychological Review, 94(2), 115–147.

Biederman, I. & Ju, G. (1988). Surface versus edge-based determinants of visual recognition. Cognitive Psychology, 21(1), 38–64.

Boynton, R. M., Fargo, L., Olson, C. X., & Smallman, H. S. (1989). Category effects in color memory. Color Research and Application, 14(5), 229–234.

Bradley, A., Switkes, E. & De Valois, K. (1988). Orientation and spatial frequency selectivity of adaptation to color and luminance gratings. Vision Research, 28(7), 841–856.

Bramao, I., Reis, A., Petersson, K. M., & Faisca, L. (2011). The role of color information on object recognition: A review and meta-analysis. Acta Psychologica, 138, 244-253.

Bramao, I., Faisca, L., Magnus, K., & Reis, A. (2012a). The contribution of color to object recognition. In Ed. I. Kypraios, Advances in Object Recognition Systems, Oxford University Press: Oxford.

Bramao, I., Francisco, A., Inácio, F., Faísca, L., Reis, A., & Petersson, K. M. (2012b). Electrophysiological evidence for colour effects on the naming of colour diagnostic and noncolour diagnostic objects. Visual Cognition, 20(10), 1164–1185.

Bruner, J. S. & Postman, L. (1949). On the perception of incongruity: a paradigm. Journal of Personality, 18, 206–223.

Callaghan, T. C. (1984). Dimensional interaction of hue and brightness in preattentive field segregation. Perception & Psychophysics, 36, 25–34.

Cavanagh, P. (1987). Reconstructing the third dimension: Interactions between color, texture, motion, binocular disparity, and shape. Computer vision, Graphics & Image Processing, 37, 171–195.

Cavanagh, P., Tyler, C. W., & Favreau, O. E. (1984). Perceived velocity of moving chromatic gratings. JOSA A, 1(8), 893–899.

Clifford, C. W. G., Spehar, B., Solomon, S. G., Martin, P. R. & Qasim, Z. (2003). Interactions between color and luminance in the perception of orientation. Journal of Vision, 3(2):1.

De Valois, K. K. & Switkes, E. (1983). Simultaneous masking interactiosn between chromatic and luminance gratings. Journal of the Optical Society of America, 73(1), 11–18.

Delk, J. L. & Fillenbaum, S. (1965). Differences in perceived color as a function of characteristic color. American Journal of Psychology, 78(2), 290–293.

Duncker, K. (1939). The influence of past experience upon perceptual properties. The American Journal of Psychology, 52(2), 255–265.

Dzhelyova, M. & Rossion, B. (2014). The effect of parametric stimulus size variation on individual face discrimination indexed by fast periodic visual stimuliation. BMC Neuroscience, 15:87, 1–12.

Elsner, A. (1978). Hue difference contours can be used in processing orientation information. Perception & Psychophysics, 24(5), 451–456.

Frome, F. S., Buck, S. L., & Boynton, R. M. (1981). Visibility of borders: separate and combined effects of color differences, luminance contrast, and luminance level. Journal of the Optical Society of America, 71(2), 145–150.

Gerth, C., Delahunt, P. B., Crognale, M. A., & Werner, J. S. (2003). Topography of the chromatic pattern-onset VEP. Journal of Vision, 3, 171–182.

Gregory, R. L. (1966). Eye and Brain: The Psychology of Seeing. McGraw Hill: NewYork.

Grossberg, S. & Mingolla, E. (1985). Neural dynamics of perceptual grouping: Textures, boundaries, and emergent segmentations. Perception & Psychophysics, 38, 141–171.

Hagmann, C. E. & Potter, M. C. (2016). Ultrafast scene detection and recognition with limited visual information. Visual Cognition, 24(1), 2–14.

Hagen, S. Vuong, Q. C., Scott, L. S., Curran, T., & Tanaka, J. W. (2014). The role of color in expert object recognition. Journal of Vision, 14(9):9, 1-13.

Hansen, T., Olkkonen, M., Walter, S. & Gegenfurtner, K. R. (2006). Memory modulates color appearance. Nature Neuroscience, 9(11), 1367–1368.

Hanslmayr, S., Pastotter, B., Bauml, K.-H. & Gruber, S. (2008). The electrophysiological dynamics of interference during the Stroop task. Journal of Cognitive Neuroscience, 20(2), 215–225

Harper, R. S. (1953). The perceptual modification of colored figures. The American Journal of Psychology, 66(1), 86–89.

von Helmholtz, H. (1867). Handbuch der Physiologischen Optik, Vol. II, published by Leopold Voss: Leipzig. In J.P.C. Southall (Ed.), Translation (Helmoltz’s Treatise on Physiological Optics, 1909), Optical Society of America: Washington, D.C., 1924, pp. 286–287.

Hering, E. (1920). Grundzüge der Lehre vom Lichtsinn. Berlin: Springer. (English version: Outlines of a Theory of the Light Sense. 1964. Translated by L.M. Hurvich and D. Jameson. Cambridge, MA: Harvard University Press).

Humphrey, G. K., Goodale, M. A., Jakobson, L. S. & Servos, P. (1994). The role of surface information in object recognition: studies of a visual form agnosic and normal subjects. Perception, 23(12), 1457–1481.

Jacques, C., Retter, T.L., Rossion, B. (2016). A single glance at a face generates larger and qualitatively different category-selective spatio-temporal signatures than other ecologically-relevant categories in the human brain. NeuroImage, 137, 21–33.

Johnson, E. N., Hawken, M. J., & Shapley, R. (2001). The spatial transformation of color in the primary visual cortex of the macaque monkey. Nature Neuroscience, 4(4), 409–416.

Johnson, E. N. & Mullen, K. T. (2016). Color in the Cortex. In Human Color Vision (pp. 189-217). Springer International Publishing.

Joseph, J. E. & Proffitt, D. R. (1996). Semantic versus perceptual influences of color in object recognition. Journal of Experimental Psychology: Learning, Memory, and Cognition, 22(2), 407–429.

Kilmer, J. M. & Lemon, R. N. (2013). What we know currently about mirror neurons. Current Biology, 23(23), R1057–62.

Kingdom, F. A. A., Beauce, C. & Hunter, L. (2004). Coulour vision brings clarity to shadows. Perception, 33(8), 907–914.

Krauskopf, J., & Farell, B. (1991). Vernier acuity: effects of chromatic content, blur and contrast. Vision Research, 31(4), 735–749.

Lee, R. J. & Mather, G. (2019). Chromatic adaptation from achromatic stimuli with implied color. Attention, Perception & Psychophysics.

Lee, R. J., Mollon, J. D., Zaidi, Q., & Smithson, H. E. (2009). Latency characteristics of the short-wavelength-sensitive cones and their associated pathways. Journal of Vision, 9(12):5, 1-17.

Liebe, S.,Fischer, E, Logothetis, N. K. & Rainer, G. (2009). Color and shape interactions in the recognition of natural scenes by human and monkey observers. Journal of Vision, 9(5):14, 1-16.

Lotto, R. B. & Purves, D. (2002). The empirical basis of color perception. Consciousness and Cognition, 11, 609–629.

Livingstone, M. S. & Hubel, D. H. (1984). Anatomy and physiology of a color system in the primate visual cortex. Journal of Neuroscience, 4(1), 309–356.

Lloyd-Jones, T. J., Roberts, M. V., Leek, E. C., Foquet, N. C., & Truchanowicz, E. G. (2012). The time course of activation of object shape and shape+colour representations during memory retrieval. PLOS One, 7(11), e48550.

Lu, A., Xu, G., Jin, H., Mo, L., Zhang, J., & Zhang, J. X. (2010). Electrophsyiological evidence for effects of color knowledge in object recognition. Neuroscience Letters, 469, 405–410.

Lupyan, G. (2015). Object knowledge changes visual appearance: Semantic effects on color afterimages. Acta Psychologica, 161, 117–130.

Markoff, J. L. (1972). Target recognition performance with chromatic and achromatic displays (Research Rep. No. SRM-148). Honeywell: Minneapolis, MN.

Marr, D. & Nishihara, H. K. (1978). Representation and recognition of the spatial organization of three-dimensional shapes. Proceedings of the Royal Society B, 200(1140).

Martin, A., Haxby, J. V., Lalonde, F. M., Wiggs, C. L., & Ungerleider, L. G. (1995). Discrete cortical regions associated with knowledge of color and knowledge of action. Science, 270(5233), 102–105.

Matkovic, K., Neumann, L., Neumann, A., Psik, T. & Purgathofer, W. (2005). Global contrast factor – a new approach to image contrast. Computational Aesthetics in Graphics, Visualization and Imaging, 159–167.

McCollough, C. (1965). Color adaptation of edge-detectors in the human visual system. Science, 149(3688), 1115–1116.

Mitterer, H. & de Ruiter, J. P. (2008). Recalibrating color categories using world knowledge. Psychological Science, 19(7), 629–634.

Moutoussis, K. (2015). The physiology and psychophysics of the color-form relationship: a review. Frontiers in Psychology, 6, 1407.

Mullen, K. T. (1985). The contrast sensitivity of human colour vision to redlJgreen and blue□yellow chromatic gratings. The Journal of physiology, 359(1), 381–400.

Nagai, J., & Yokosawa, K. (2003). What regulates the surface color effect tin object recognition: Color diagnosticity or category? Technical Report on Attention and Cognition, 28, 1–4.

Naor-Raz, G., Tarr, M. J., & Kersten, D. (2003). Is color an intrinsic property of object representation? Perception, 32, 667–680.

Norcia, A. M., Appelbaum, L. G., Ales, J. M., Cottereau, B. R. & Rossion, B. (2015). The stead-state visual evoked potential in vision research: A review. Journal of Vision, 15(6):4, 1-46.

Nunez, V., Shapley, R. M. & Gordon, J. (2007). Nonlinear dynamics of cortical responses to color in the human cVEP. Journal of Vision, 17(11):9, 1-13.

Olkkonen, M., Hansen, T. & Gegenfurtner, K. R. (2008). Color appearance of familiar objects: Effects of object shape, texture, and illumination changes. Journal of Vision, 8(5):13, 1-16.

Oliva, A. & Torralba, A. (2001). Modeling the shape of the scene: a holistic representation of the spatial envelope. International Journal of Computer Vision, 42(3), 145–175.

Oostenveld, R. & Praamstra, P. (2001). The five percent electrode system for high-resolution EEG and ERP measurements. Clinical Neurophysiology,112(4), 713–719.

Or, C.-F., Retter, T. L. & Rossion, B. (2019). The contribution of color information to rapid face categorization in natural scenes. Journal of Vision, 19(5), 1–20.

Pinna, B., Brelstaff, G. & Spillmann, L. (2001). Surface color from boundaries: a new ‘watercolor’ illusion. Vision Research, 41, 2669–2676.

Price, C. J. & Humphreys, G. W. (1987). The effects of surface detail on object categorization and naming. The Quarterly Journal of Experimental Psychology, 41A(4), 797-828.

Proverbio, A. M., Burco, F., del Zotto, M., & Zani, A. (2004). Blue piglets? Electophysiological evidence for the primacy of shape over color in object recognition. Cognitive Brain Research, 18, 288–300.

Quek, G. L. & Rossion, B. (2017). Category-selective human brain processes elicited in fast periodic visual stimulation streams are immune to temporal predictability. Neuropsychologia, 104, 182–200.

Rabin, J., Switkes, E., Crognale, M. A., Schneck, M. E. & Adams, A. J. (1994). Visual evoked potentials in three-dimensional color space: Correlates of spatio-chromatic processing. Vision Research, 34, 2657–2671.

Regan D. (1966). Some characteristics of average steady-state and transient responses evoked by modulated light. Electroencephalography and Clinical Neurophysiology, 20 (3), 238–248.

Retter, T. L. & Rossion, B. (2016). Uncovering the neural magnitude and spatio-temporal dynamics of natural image categorization in a fast visual stream. Neuropsychologia, 91, 9–28.

Retter, T. L., Gwinn, O. S., O’Neil, S., Jiang, F., & Webster, M. A. (2020). Neural correlates of perceptual color inferences as revealed by #thedress. Journal of Vision, 20(3):7, 1-20.

Riggs, L. A. & Sternheim, C. E. (1968). Human retinal and occipital potentials evoked by changes of the wavelength of the stimulation light. Journal of the Optical Society of America, 59(5), 635–640.

Robson, A. G. & Kulikowski, J. J. (1998). Objective specification of tritanopic confusion lines using visual eboked potentials. Vision Research, 38, 3241–3245.

Rossion, B. & Pourtois, G. (2004). Revisiting Snodgrass and Vanderwart’s object pictorial set: The role of surface detail in basic-level object recognition. Perception, 33, 217–236.

Rossion, B., Prieto, E. A., Boremanse, A., Kuefner, D. & Van Belle, G. (2012). A steady-state visual evoked potential approach to individual face perception: effect of inversion, contrast-reversal and temporal dynamics. NeuroImage, 63, 1585–1600.

Rossion, B., Torfs, K., Jacques, C. & Liu-Shuang, J. (2015). Fast periodic presentation of natural face images reveals a robust face-selective electrophysiological response in the human brain. Journal of Vision, 15(1):18, 1-18.

Seymour, K., Clifford, C. W., Logothetis, N. K., & Bartels, A. (2010). Coding and binding of color and form in visual cortex. Cerebral Cortex, 20(8), 1946–1954.

Shevell, S. K. and Kingdom, F. A. A. (2008). Color in complex scenes. Annual Review of Psychology, 59, 143–166.

Slotnick, S. D. (2009). Memory for color reactivates color processing region. Cognitive Neuroscience and Neurophysiology, 20, 1568–1571.

Tanaka, J. W. & Presnell, L. M. (1999). Color diagnosticity in object recognition. Perception & Psychophysics. 61(6), 1140–1153.

Tanaka, J. Weiskopf, D. & Williams, P. (2001). The role of color in high-level vision. Trends in Cognitive Science. 5(5), 211–215.

Therriault, D. J., Yaxley, R. H., & Zwaan, R. A. (2009). The role of color diagnosticity in object recognition and representation. Cognitive Processing, 10(4), 335–342.

Thompson, E. (1995). Colour vision, evolution, and perceptual content. Synthese, 104, 1–32.

Thorell, L. G., De Valois, R. L., & Albrecht, D. G. (1984). Spatial mapping of monkey VI cells with pure color and luminance stimuli. Vision research, 24(7), 751–769.

Tiechmann, L., Grootswagers, T., Carlson, T., & Rich, A. N. (2018). Seeing versus knowing: the temporal dynamics of real and implied colour processing in the human brain. BioX archive.

Tiechmann, L., Quek, G. L., Robinson, A., Grootswagers, T., Carlson, T., & Rich, A. N. (2019). Yellow strawberries and red bananas: The influence of object-colour knowledge on emerging object representations in the human brain. BioX archive.

Triesman, A. (1982). Perceptual grouping and attention in visual search for features and for objects. Journal of Experimental Psychology: Human Perception and Performance, 8(2), 194–214.

Trosianko, T. & Harris, J. P. (1988). Phase discrimination in compound chromatic gratings. Vision Research, 28, 1041–1049.

Uchikawa, K., & Shinoda, H. (1996). Influence of basic color categories on color memory discrimination. Color Research and Applications, 21(6), 430–439.

Van Lier, R., Vergeer, M., & Anstis, S. (2009). Filling-in afterimage colors between the lines. Current Biology, 19(8), R323–R324.

Van Gulick, A., & Tarr, M. (2010). Is object color memory categorical? Journal of Vision, 10(7):407, 407a.

Vandenbroucke, A. R. E., Fahrenfort, J. J., Meuwese, J. D. I., Scholte, H. S., & Lamme, V. A. F. (2016). Prior knowledge about objects determines neural color representation in human visual cortex. Cerebral Cortex, 26, 1401–1408.

Walls, G. L. (1942). The vertebrate eye and its adaptive radiation. Cranbook Instittue of Science: Bloomfields Hills, MI.

Webster, W. A., De Valois, K. K. & Switkes, E. (1990). Orientation and spatial-frequency discrimination for luminance and chromatic gratings. Journal of the Optical Society of America A, 7(6), 1034–1049.

Winkler, A., Spillmann, L., Werner, J. S. & Webster, M. A. (2015). Asymmetries in blue-yellow color perception and in the color of “the dress”. Current Biology, 25(13), R547–R548.

Witzel, C., Valkova, H., Hansen, T., & Gegenfurtner, K. R. (2011). Object knowledge modulates colour appearance. i-Perception, 2, 13-50. dx.doi.org/10.1068/i0396

Wolfe, J. M., Cave, K. R. & Franzel, S. L. (1989). Guided search: an alternative to the feature integration model for visual search. Journal of Experimental Psychology: Human Perception and Performance, 15(3), 419–433.

Wolfe, J. M. & Horowitz, T. S. (2004). What attributes guide the deployment of visual attention and how do they do it? Nature Reviews Neuroscience, 5, 1–7.

Wurm, L. H., Legge, G. E., Isenberg, L. M., & Luebker, A. (1993). Color improves object recognition in normal and low vision. Journal of Experimental Psychology: Human Perception and Performance, 19(4), 899–911.

Yao, A. Y. J., & Einhauser, W. (2008). Color aids late but not early stages of rapid natural scene recognition. Journal of Vision, 8(16):12, 1–13,

Zeki, S., & Shipp, S. (1988). The functional logic of cortical connections. Nature, 335(6188), 311–317.

